# All protein aggregates will thermodynamically order

**DOI:** 10.1101/2020.09.15.278408

**Authors:** Aleksandra W. Nielsen, Levent Sari, Rowan Fraser, Milo M. Lin

**Author notes:** These authors contributed equally (Aleksandra W. Nielsen, Levent Sari).

## Abstract

Proteins can aggregate into disordered liquid droplets or ordered assemblies such as amyloid fibrils. These two distinct phases determine the spatial organization within cells and serve differing roles in a wide range of biological functions including gene regulation, organelle and synapse formation, and memory consolidation. The ordered phase can also give rise to diseases such as Alzheimer’s. However, how the protein sequence determines aggregation fate is an open question. Here we establish a general statistical mechanical theory of the disordered-to-ordered transition for polymer aggregates, including proteins, thereby completing the phase diagram for this general class of matter. The theory produces a simple universal equation determining the favored phase as a function of the temperature, polymer length, and inter-residue interaction energy variance. We show that the sequence-dependent energy variance can be efficiently calculated from all-atom molecular dynamics simulations, so that the theory has no adjustable parameters. The equation accurately predicts the experimental length-dependent crystallization temperature of synthetic polymers. The theory shows that all protein aggregates, regardless of sequence, will thermodynamically order, even in the most extreme thermophiles. Therefore, energy must be expended to maintain the disordered protein aggregate at steady state. More broadly, the theory establishes a lower bound on the ordering transition temperature for polymers. This bound indicates that condensates of any organic polymer will spontaneously order on all habitable planets.

---

Polymers in solution can exist in dilute, disordered aggregate, or ordered aggregate states corresponding to gas, liquid, and solid phases, respectively. Proteins *in vivo* can manifest in all of these phases, with each phase giving rise to fundamentally different functional and toxicity properties. Proteins typically function in the dilute phase as individual polymers or small complexes of a few polymers. Ordered aggregates such as cytoskeletal and amyloid fibrils effect global decisions such as programmed cell death^1^, cell-cycle progression^2^, and the consolidation of long-term memory^3^. Such aggregation is sensitively tuned by a host of modulatory proteins in response to changing conditions^4–6^. Ordered aggregates in the form of amyloid fibrils are also implicated in a wide range of neurodegenerative diseases^7–11^. Once formed, some amyloids can become prions, which are preserved after cell division^8^ and exported to convert new proteins^12^ as a form of structure-based inheritance. Finally, proteins also aggregate into the disordered phase as membraneless organelles such as nucleoli and P granules^13^. Recently, the ability to regulate cell function by controlling the formation and segregation of these liquid phases has been reported in a wide range of functions, including noise reduction^14^, chromatin regulation^15^, organelle formation^16^, and synapse remodeling^17^.

The transition from the dilute to the disordered aggregate is well understood. The Flory-Huggins theory^18,19^, which takes into account the thermodynamic principle of balancing the entropy of mixing against the mean stabilization energy of intra-polymer interactions, accurately predicts experimental aggregation transition of polymers, including proteins^20^. Sequence characteristics such as multivalency^21^ and low-complexity domains^22^ have also been demonstrated *in vitro* to tune the transition in favor of aggregation. For the direct assembly of fibrils from the dilute phase, distinct fibril morphologies can form via competing pathways^23^, and an elegant kinetic model has analytically revealed general rate-limiting mechanisms^24^.

In contrast, the disorder-order transition of protein aggregates is poorly understood. Some protein aggregates have been observed to transition from the disordered phase to ordered amyloids, although the sequence determinants allowing for this transition are unclear^25^. Because the timescale of conformational reorganization is highly heterogeneous, the transition kinetics may not always be experimentally accessible. For example, the stoichiometry of distinct fibril strains corresponding to different conformations of certain proteins does not equilibrate (i.e. preserves its preparation history) within weeks^26^. Experimentally, it has been observed that many low-concentration proteins and non-biological protein sequences can be induced to form amyloids *in vitro*^27^, leading to the hypothesis that general principles may favor the stability of the ordered phase^27,28^. The inconclusive experimental picture is underscored by the lack of a thermodynamic theory that predicts the disorder-order transition of the condensed phase as a function of polymer properties. In contrast to the Flory-Huggins model, such a theory would need to consider the conformational entropy of the disordered phase as well as interaction characteristics beyond the mean energy.

Here, we derive a statistical mechanical theory of the disorder-order transition, and show that the stability condition is determined by a simple universal phase equation that depends on three parameters: polymer length, interaction energy variance, and temperature. We demonstrate that the interaction energy distribution is efficiently and accurately calculated using all-atom molecular dynamics simulations. Consequently, we use the phase equation to predict the length-dependent experimental crystallization temperature of a variety of synthetic polymers with no adjustable parameters. For proteins, the phase equation establishes that aggregates of all possible protein sequences, over all physiologically realizable temperatures, will thermodynamically order. The theory also shows that the minimum complexity sequence, alkane, sets the lower bound on the order-disorder transition temperature for all polymers. Because the alkane transition temperature is close to the maximum temperature for habitable planets, we conclude that the tendency to spontaneously order is a generic driving force for any polymer-based life.

## Disorder-order transition theory for polymer condensates

Consider the disordered aggregate composed of *N* copies of a polymer (e.g. protein) of length *l. N* is typically so large that the system approaches the thermodynamic limit in which the intensive variables such as the average structure is independent of extensive variables such as *N. l* is typically between 10 and 1000. To characterize the energy of an aggregate, we first consider the distribution of interaction energies of all pairs of residues (e.g. amino acids). Let *p*(*ϵ*) be the probability that the interaction energy of a residue with its neighbors in the aggregate has energy *ϵ*. Define the mean and standard deviation of *ϵ* to be 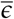 and *σ*, respectively. Then, the total energy, *E*, of an aggregate microstate is the sum of 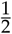*Nl* such interactions, where the factor of 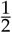 is due to double-counting. The probability distribution of *E, P*(*E*), is therefore the 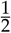 *Nl*’th convolution of *p*(*ϵ*). Because the polymer constraint enforces the concentration of each type of residue to be fixed over length scales much larger than the polymer length *l*, the residue interactions of the disordered phase are well-mixed. Therefore, regardless of the shape of *p*(*ϵ*), *P*(*E*) approaches a normal distribution by the Central Limit theorem because *Nl* is enormous: 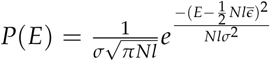. The free energy of the set of microstates with energy *E* is: *G*(*E*) = *E − TS*(*E*), where the entropy of the set of microstates of energy *E, S*(*E*), is equal to Boltzmann’s constant *k* times the logarithm of the number of conformational microstates of the aggregate with energy between *E* and *E* + *dE*. The latter is equal to the total number of possible microstates, which we define to be Ω, times *P*(*E*)*dE*. Therefore,

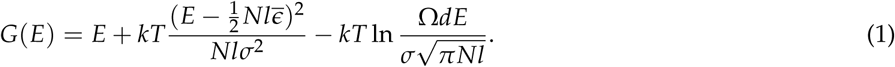

In the thermodynamic limit of vanishing *dE*, its value does not affect the difference in free energy between states, which is the relevant thermodynamic quantity. From Eq. 1, the free energy is seen to be a quadratic function of *E*, where *E* goes from *−*∞ to ∞; however, the physically possible range of *E* is bounded by the minimum and maximum possible energy. Any energy value outside of this bound corresponds to fewer than one microstate: these energy values are probabilistically prohibited because they have “run out of entropy.” We identify the unique single microstate whose energy is the minimum of *E* as the ground state ordered phase. Therefore, whether the ordered phase is more stable than the disordered phase depends on whether the minimum of *G*(*E*) is found within this energy range. If the derivative of *G*(*E*) with respect to *E* at the minimum energy is negative, then the lowest free energy macrostate is a high entropy (disordered) state (black square in Fig. 1b). Otherwise, the minimum energy ordered state corresponds to the free energy minimum (orange square in Fig. 1b). By solving for this criterion at the lowest energy state (*S*(*E*_min_) = 0), we obtain the condition under which the ordered state is thermodynamically favored (See Supp. information):

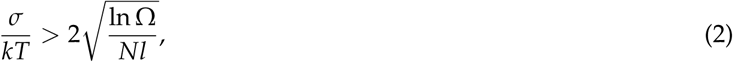

which occurs when the heterogeneity (*σ*) of residue interactions (normalized by *kT*) exceeds the square root of the entropy (ln Ω) of polymer configurations (normalized by the total number of residues *Nl*).

**Figure 1:**
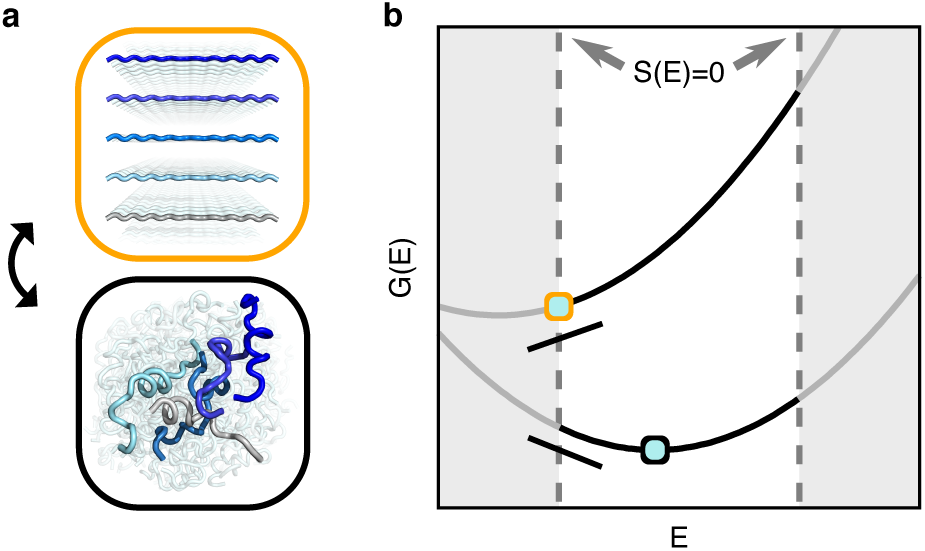
Statistical mechanical model of the order-disorder transition. The ordered (orange square) or disordered (black square) phase is stable (a), depending on the minimum free energy state as a function of energy (b). Representative curves in (b) correspond to systems whose free energy derivative at the lowest energy state are positive and negative, corresponding to thermodynamic stability of the ordered versus disordered phases, respectively.

Unlike the condition for aggregation set out by Flory-Huggins theory, the condition for ordering is not dependent on the mean interaction energy, but rather the variance. This is because the interaction density is preserved between the ordered and disordered phases. To obtain Ω, we derived the conformational entropy of *N* chains of length *l* in a self-avoiding dense aggregate (See Supplementary information). The final phase equation, which sets the condition for thermodynamic stability of the ordered phase, is:

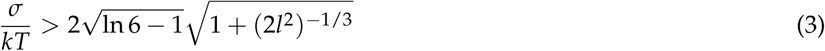

and is independent of the extensive variable *N*, as expected. The order-disorder transition temperature is thus weakly monotonically increasing with chain length, and asymptotes in the long chain limit to:

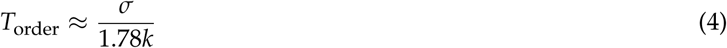

This result is simply modified to include the entropy of mixing for the order-disorder transition of mixtures of different types of polymers (See Supplementary information).

## Theory predicts length-dependent experimental and simulation data

To test the theory, we chose the well-characterized alkane and nylon-6 polymers for two reasons: (i) unlike polypeptides (proteins), their length-dependent melting temperatures have been measured^29–31^ and (ii) along with polypeptides, their backbones are constructed solely from the set of carbon-carbon and carbon-nitrogen (amide) bonds, with nylon being a chemical interpolation between alkanes and polypeptides (Fig. 2a). After quantifying the accuracy of Eq. 3 on these polymers, we can then use it to make predictions for proteins of varying lengths and sequence compositions (See next section).

**Figure 2:**
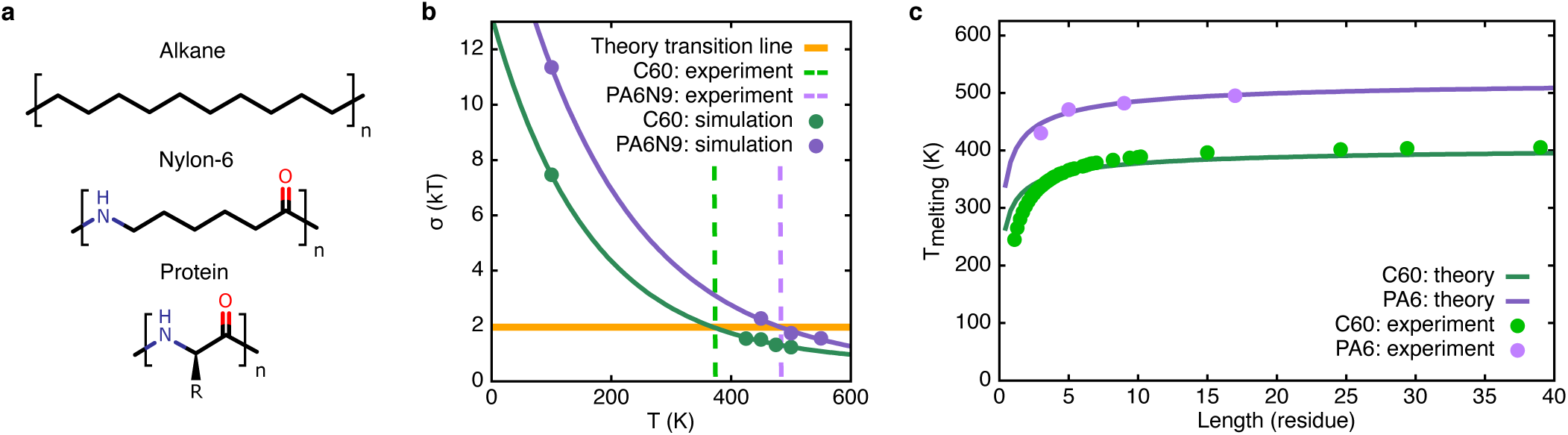
Parameter-free prediction of experimental data. a, Chemical structures of polymers for which σ is calculated. The size of a residue, which is based the calculated polymer-dependent Kuhn length, is enclosed in square brackets (See Text). b, The predicted theoretical melting temperatures for C60 (alkane) and PA6N9 (nylon-6) correspond to the intersection of σ/kT with the theoretical ordering condition (Eq. 3; solid orange line for both C60 and PA6N9 due the two polymers having similar l). Experimental melting temperatures are shown as dashed lines^29,30^. c, Length dependence of transition temperatures, with dots representing experimental data^29–31^ and solid lines theoretical curves.

To predict the transition (melting) temperature, we performed MD simulations to calculate *σ* at a range of temperatures and found the intersection of the interpolation of these points with the theoretical transition condition Eq. 3 (Fig. 2b). We tested three of the most-used force field parameter sets: AMBER99SB^32^, Gromos54a-7^33^, and L-OPLS-AA^34^. These force fields produced small variations in *σ*. Because AMBER99SB gave the median value (Fig. S2), all simulation data, unless otherwise noted, are produced using this force field. The results for hexacontane (C60) and 9-mer nylon-6 polymer (PA6N9), along with the experimental melting temperatures, are shown in Fig. 2b. Because of the slight temperature dependence of the *σ* values, simulations were repeated at several different temperatures to be able to interpolate the data and subsequently estimate *σ* at the exact order-disorder transition line. As a result, the predicted melting temperatures of 367 and 485 K are in very good agreement with the experimental values of 372^29^ and 482 K^30^ for C60 alkane and PA6N9 nylon-6, respectively.

We also calculated the length dependence of melting temperatures using Eq. 3. For each polymer, the residue length is chosen to be equal to the calculated Kuhn length to be consistent with the length units used in the theory (See Supplementary materials for details). Fig. 2c shows the excellent agreement with the experimental data for both alkanes and nylon-6^29–31^. In addition to the overall quantitative agreement, the strong length dependency for short polymers and the saturation of length dependence for long polymers are consistent characteristics of theory and experiment.

We noted that, of all the polymers studied, only alkanes could make the disorder-order transition within the time window of all-atomic MD simulations (sub-microseconds) by spontaneous formation of crystalline microdomains (See S3). This means that if using different force-field parameter sets leads slightly different *σ* values for alkanes, then the crystallization temperature observed using MD should also vary according to Eq. 3. This would be an orthogonal way to test the theory using only simulation data. Therefore, we all C60 simulations using 3 other force-fields: Gromos54a-7^33^, L-OPLS-AA^34^, and TraPPE-UA^35^. The first two of these, like AMBER99SB, are at all-atom resolution, while the last one (TraPPE-UA) is a coarse grained force-field for which hydrogen atoms are united with their corresponding heavy atoms. The versus observed melting temperatures for these four force-fields are shown in Fig. 3 for C60. The predictions using the all-atom force-fields are in close proximity to the experimental value of 372 K. On the other hand, the prediction of the coarse-grained force-field, 230 K, deviates significantly. Nevertheless, Eq. 3 accurately predicts the observed crystallization temperature corresponding to each of the four force fields using the *σ* calculated from the disordered phase (Fig. 3). In contrast, the crystallization temperature is not correlated with the mean interaction energy, with the sign of the correlation changing depending on whether the coarse-grained force-field is included or not (Fig. 3 inset). As dictated by the theory, the *variance* in interaction energy, not the mean, determines the transition temperature.

**Figure 3:**
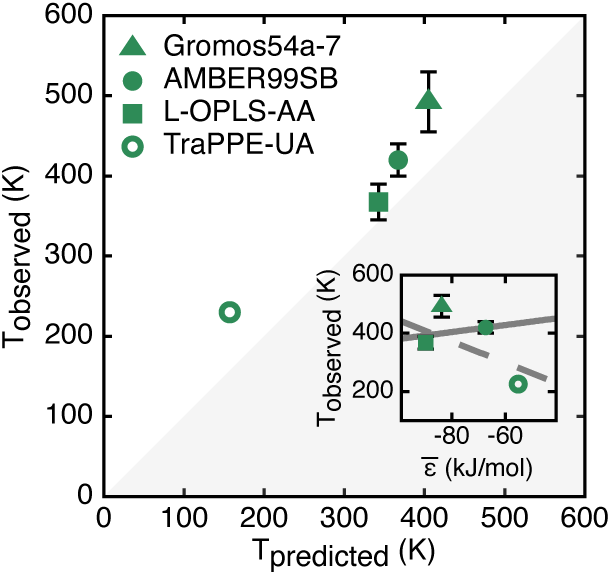
Testing the theory using force-field variability. The C60 crys- tallization temperatures observed us- ing different force fields are compared to those predicted by Eq. 3 using the calculated σ. The observed temperature is not correlated with the mean interaction energy (inset), including (dashed line) or excluding (solid line) the united-atom force-field.

## All protein condensates are ordered in the thermodynamic limit

Next, we used Eq. 3 to predict the disorder-order transition for polypeptides using the same protocol as for the synthetic polymers. We obtained the interaction energy distribution for each amino acid type by performing MD simulations of 100 different proteins from the PDB database (Table S1) and sampling the interactions within the protein cores for 100 nanoseconds each (See Supplementary data). The resultant energy distributions are given in Fig. 4a for all twenty amino acids. Energy distributions for hydrophobic residues are narrower than those of polar and charged residues. The widest distributions were observed for the four charged amino acids (D, E, R, K), consistent with their diverse chemical interactions involving hydrogen bonding, pure electrostatic interactions, and salt bridges. Because *σ* for all individual amino acids exceed the ordering threshold set by Eq. 3, this already indicates that *σ* for polypeptide chains will also exceed the threshold. As homopolymers likely represent a lower bound on *σ*, we first showed that *σ* values of all aggregating homo-polypeptides, including polyglycine, are well above the transition line, even at *T* = 400*K* (Fig. 4b). This suggests that all protein sequences are also above the transition line.

**Figure 4:**
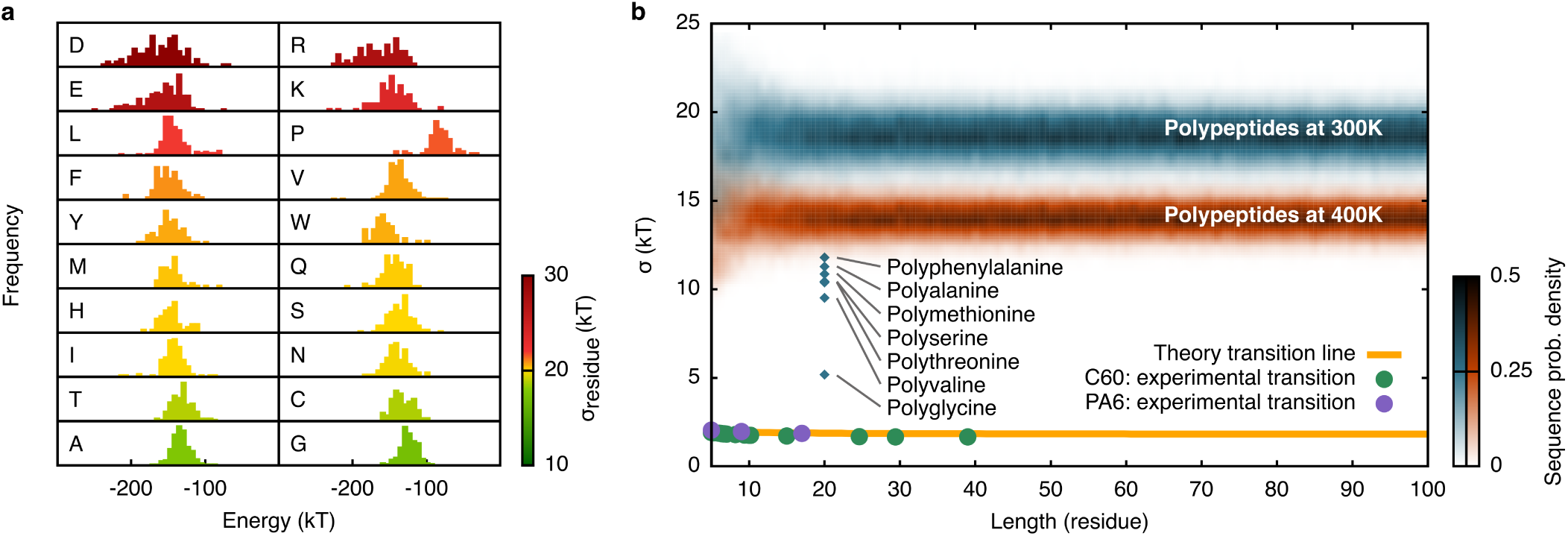
Order-disorder transition over all polypeptide lengths and sequences. a, Distributions of interaction energies for 20 amino acids with their environments, as calculated over 100 different protein simulations. b, Standard deviations for a total of 100,000 randomly generated polypeptide sequences (1,000 for each length), based on the interaction energies given in a. σ values at experimental melting temperatures for alkanes and nylon6 are also shown along the predicted transition line, as given by Eq. 3. Blue diamonds represent homo-polypeptides simulated explicitly.

In order to calculate *σ* for a large representative population of heteropolypeptides, explicit simulation of each polypeptide would be infeasible. Instead, we employed an additive model in which *σ* for a hetero-polypetide is calculated using the energy distribution of the weighted superposition of the amino acid energy distributions (Fig. 4a) of the residues composing the sequence (See Supp. methods). This additive approach yields similar *σ* values (within a few *kT*) as explicit bulk-phase simulation of hetero-polypeptides (Fig. S5). We generated 1,000 random sequences for each polypeptide length (a total of 100,000 sequences from *L* = 5 to *L* = 104). The standard deviation of the resultant distribution for each sequence are given in Fig. 4b at both 300 and 400 K, with the latter representing the upper temperature limit for life on Earth^36^. All sequences are well above the transition line, indicating that the ordered aggregate is the thermodynamically stable state.

## Discussion

Up to now, the phase diagram of polymers such as proteins was incomplete because the rules for the stability of the ordered phase (e.g. amyloid) relative to the disordered liquid phase were unknown. In this study, we establish a statistical mechanical theory for this transition, which culminates in a simple phase equation predicting the stability as a function of polymer length, variance of monomer-interaction energy *σ*^2^, and temperature. By performing atomistic molecular dynamics simulations on the disordered state, *σ* can be quickly and accurately calculated for any polymer. For polymer aggregates whose phase behavior have been calorimetrically measured, the phase equation accurately predicts experimental order-disorder transition temperatures in a length-dependent manner, with no fitting parameters. The phase equation establishes that all protein aggregates, for all sequences and feasible temperatures (i.e. below the 500K biomolecular stability limit^37^), are thermodynamically stable in the ordered phase. These results substantiate a long-standing conjecture by Astbury^28^ and, later, Dobson et al^38^, about the ubiquitous nature of ordered protein aggregates such as amyloids. Using the key insight that the disorder-order transition is determined by *σ*, we can generalize this conclusion further to all organic polymers. Because the transition temperature of the simplest organic polymer, alkane, is near 400K, we infer that polymer-based aggregation will spontaneously order in the thermodynamic limit, even if composed of alternative chemistries on one of the catalogued habitable exoplanets, whose surface temperature range is defined to be below 395 K^39^.

A universal thermodynamic tendency to order implies that all proteins must be regulated in terms of concentration, kinetic barriers and/or by active (i.e. energy-consuming) agents such as chaperones if they are to avoid forming ordered aggregates such as amyloids. Conversely, the universal tendency to order also serves as a spontaneous self-organizational principle that can facilitate biochemical information storage and transfer. For example, prion-based inheritance has been proposed as a simple auto-catalyzed reaction that could facilitate the pre-biotic to biotic transition^40^. The ability of existing prion proteins to switch the cellular state^41^, and to facilitate trans-kingdom metabolic signaling within mixed microbial communities^42^ support this hypothesis. This work suggests that this mechanism of self-organized information transfer is not only possible, but omnipresent.

## Materials and methods

The molecular dynamics simulations (MD) were performed using GROMACS 5.0.4^43^ software. For protein simulations, we used 100 different protein structures (Table S1) with SPCE^44^ explicit water molecules, employing AMBER99SB-ildn^45^ force field. For each protein structure, 100 ns long full atomistic MD trajectories were obtained (10 microseconds in total). In post-simulation analyses, solvent exposed mobile residues were excluded, concentrating only on the inner cores of each protein structure to mimic bulk medium. The interaction energy of each residue with all other residues within 1 nm proximity was calculated, and energy distributions for all 20 amino acids were obtained by taking average values over each trajectory. These amino acid energy distributions were used to calculate overall *σ* of the randomly generated polypeptide sequences.

For bulk medium simulations, we performed atomistic molecular dynamics simulations of C60 alkane, and repeated the simulations using 4 different force-fields: Gromos54a7^33^, AMBER99SB^32^, L-OPLS^34^ and TraPPE-UA.^35^ For PA6N9 nylon-6 and polypeptide systems, we used AMBER99SB and AMBER99SB-ildn^45^ force fields, respectively. Prepared boxes consisted of 2,376 C60, 2,772 PA6N9 and 2,704 polypeptide (boxes 1-6 in Table S2, as well as homo-polypeptides in Fig. 4b) chains. For polypeptides, additional systems were created, with 2,000 chains put randomly in space (boxes 7-14 Table S2). First, initially prepared boxes were mixed at high temperatures, and then the resulting aggregates were quenched and equilibrated to different target temperatures, after which production simulations were performed to calculate *σ*. Mean interaction energies were calculated between a residue (defined as Kuhn length) and all atoms within 1 nm proximity, and then averaged over a time window during which a microstate does not change (Fig. S1). The interaction energies of at least 600 different residues were used for standard deviation prediction. In case of heterogenous chains, we superposed mean energies of different residues to obtain the overall *σ*. To observe the crystallization temperature of C60, 25 ns simulations were performed for all explicit-atom force fields at increasing temperatures. The lower bound of the crystallization temperature is taken to be the highest temperature at which spontaneous ordering was observed while its upper bound was determined to be the lowest temperature in which the nucleated ordered domains melted. Because spontaneous ordering was not observed for the united-atom force field within 25 ns, a bipartite order-disorder system was initiated, and the ordering temperature was taken to be the temperature at which the rate of change of intensive parameters (such as density) changed sign (Fig. S4). Further MD set-up details are given in supplementary information section III, while supplementary section IV has the details in post-simulation analyses.

## Acknowledgments

The authors would like to thank Prashant Mishra, Kimberly Reynolds, Lukasz Joachimiak, and Bryan Ryder for feedback that significantly improved the manuscript. This work was supported by the Cecil and Ida Green Foundation, the Leland Fikes Foundation, and the Welch Foundation (Grant No. I-1958-20180324).

## Author contributions

M.M.L. conceived the project, developed the theoretical model, and supervised the project. A.W.N., L.S., and R.F. performed simulations. L.S. and A.W.N. analyzed simulation and experimental data. A.W.N., L.S., and M.M.L. wrote the paper.

## SUPPLEMENTARY INFORMATION

### I. Detailed derivation of order-disorder transition equation

From Eq. 1, the aggregate will be ordered if:

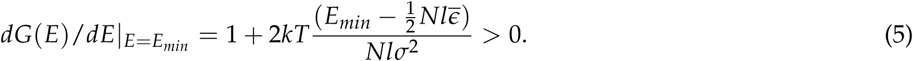

To find the lowest energy state, we note that it is the state in which the entropy is zero: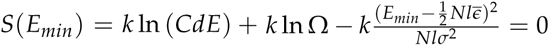. Since the first term of this expression is logarithmic in *N* whereas the others are polynomial in *N*, we can ignore the first term to obtain:

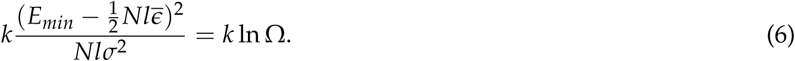

Substituting the negative root solution into the ordered-aggregate condition, we obtain the ordering condition Eq. 2:

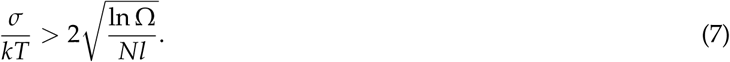

Previous work^46^ showed that the number of compact conformations of a single chain of length *L* is:

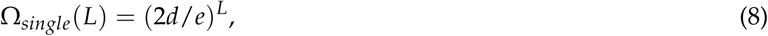

where *d* and *e* are the dimension and Euler’s constant, respectively. We would like to modify this result to calculate the number of ways to fold *N* separate chains, each of length *l*. Imagine a particular microstate consisting of all chains packed together. Starting from a randomly chosen initial chain, find the closest end of another unlinked chain and draw a linker connecting the two ends. Continue in this way until all of the chains are linked together. The typical length of a linker, *l*_*\**_, is set by the shortest distance between the endpoints of the chains:

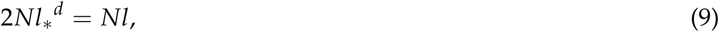

where *Nl* is the total protein volume, and there are 2*N* boxes of dimension *l*_*\**_^*d*^ that comprise the volume.Thus,

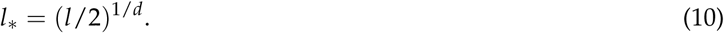

Define this procedure to be a one-to-one mapping between any conformation of *N* protein chains of length *l* and an equivalent system in which there is a single chain of length *N*(*l* + *l*_*\**_). Then we can estimate Ω:

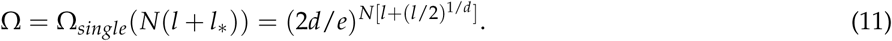

Substituting Eq. 14 into Eq. 10, we obtain the final condition in terms of the protein composition:

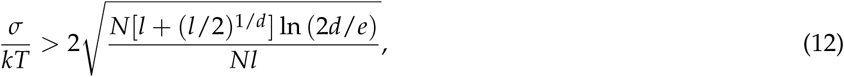

which simplifies to:

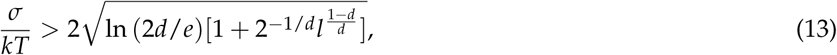

In 3 dimensional space,

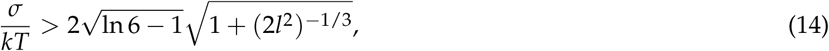

and is independent of the total number of chains, as expected. The critical variation of interaction energy, *σ*_crit_, is also weakly dependent on chain length *l*:

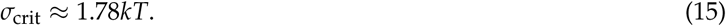

### II. Protein mixtures

The order-disorder transition will be modified if, instead of a single protein species, there are *n* different proteins. For simplicity, assume that the amino acids for all the proteins are drawn from the same distribution and that each protein is of the same length. These assumptions capture the essential nature of the effect of mixing proteins on the order-disorder transition, which is dependent on the number and stoichiometries of different protein types. If the molar fractions of each species *i* is denoted by *f*_*i*_, then the additional entropic contribution to the disordered state is:

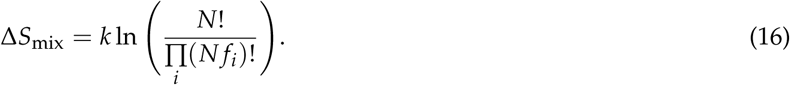

In the large *N* limit, this simplifies to:

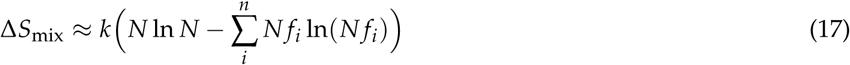

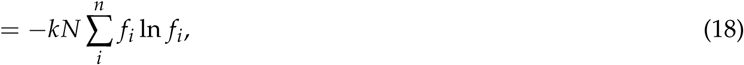

which is the Gibbs-Shannon entropy of protein identities in the mixture. The inequality for protein mixtures is thus:

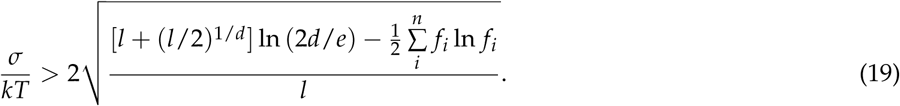

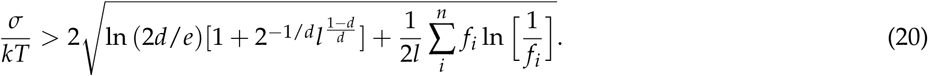

The disorder-order transition of a mixed polymer aggregate is therefore modified by the last term in the square root. Because this term is at most 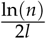, the mixing entropy has a negligible effect in the case of long polymers (large *l*).

### III. Molecular dynamics simulations

All simulations and calculations have been done on UTSW BioHPC computing clusters. Molecular dynamics simulations were performed using GROMACS 5.0.4 software^43^. For protein core simulations, we have chosen 100 different protein structures from protein data bank (see Table S1 in supplementary data for the pdb ids), and performed 100 ns long full atomistic molecular dynamics simulations on each protein. Dodecahedron box was constructed with periodic boundary conditions for each protein structure. After an initial steepest descent energy minimization, we have performed thermal equilibration where 10 ns NVT, 10 ns NPT with Berendsen^47^ barostat, and 10 ns NPT with Parrinello-Rahman^48^ barostat were employed sequentially. Hydrogen-only virtual sites^49^ with 5 fs time step, Particle Mesh Ewald (PME)^50^ summation for long-range electrostatics, and NPT ensemble were used in production level calculations. All simulation boxes have SPCE^44^ explicit water molecules, with a minimum distance of 1 nm between solute and box edge. Appropriate number of neutralizing ions (either Na or Cl) were added to each box to ensure that total charge becomes zero. AMBER99sb-ildn force-field^45^ was chosen as it is known to describe protein structures accurately^51^.

For bulk media molecular dynamics simulations of C60 alkanes, we have used several different force fields: all atomic Gromos54a7^33^, AMBER99SB^32^ and L-OPLS^34^, with an addition of united-atom TraPPE^35^. For all further polymer simulations, only specific AMBER99 force fields were employed. Triclinic boxes with initially ordered polymers were set up to allow for at least two times the length of a polymer in x, y and z directions, and therefore the number of chains per box varied: they were 2,376 C60, 2,772 PA6N9 and 2,704 polypeptide (boxes 1-6 in Table S2, as well as homo-polypeptides in Fig. 4b) chains. For polypeptides, additional systems were created, with 2,000 chains, 400 of each kind (boxes 7-14 Table S2), put randomly in space. All systems were minimized using steepest descent algorithm and then simulated under periodic boundary conditions, with either 1 or 2 fs time step. Noose-Hoover^52^ thermostat and Parrinello-Rahman^48^ barostat were used for the production phase simulations. In case of alkane simulations, initial ordered box with C60 chains was heated to 550 K and then mixed for 200 ns under NVT conditions using Gromos54a7 force field. This produced a disordered aggregate structure that we used as a starting point for further all-atom simulations. Each simulation was performed for 25 ns at 500 K and continued at lower temperatures until a spontaneous ordering was observed. Moreover, we simulated the C60 boxes at 100 K for 25 ns, trapping them in disordered aggregates. The TraPPE united-atom force field was used to run 20 ns NPT simulations every 25 K from 500 K to 200 K, with an additional one at 100 K. For PA6N9 and polypeptide bulk simulations, we have chosen to use AMBER99SB and AMBER99SB-ildn force fields, respectively. The initial ordered structure of PA6N9 was first heated under NVT and later NPT conditions at 550 K for 50 ns in total, and then equilibrated at lower temperatures (as seen in Fig. 2b). PA6N9 was also simulated at 100 K. All boxes containing polypeptide chains were simulated under NPT conditions for 25 ns at 600 K, and then for another 25 ns at 300 K.

### IV. Simulation analysis

#### Calculating *σ* from MD simulations

In order to calculate *σ* in bulk media to plug into the theory, we need to translate atomistic simulation data to the polymer model of self-avoiding random walk on a 3 dimensional lattice. The number of backbone atoms per residue of the lattice model can be obtained by computing the radius of gyration of the random walk, *R*_*g*_:

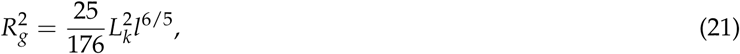

where *L*_*k*_ is Kuhn length and *l* is the number of residues in a chain^53^. *σ* calculations were based on short-range Coulomb and Lennard-Jones interaction energies between a defined residue and its surroundings within 1 nm radius. Total length of a polymer was defined by number of its Kuhn units (residues), and calculated as in Eq. 21. For polypeptides, this corresponds with the definition of a residue being one amino acid long. Standard deviations were computed from mean interaction energies of at least 600 non-terminal residues and averaged over a time window within which the configuration of the residue and its interaction neighbors would map to the same lattice microstate (Fig. S1). In the case of simulations at lower temperatures, the rearrangement time is longer than the simulation time. In this case, the calculated *σ* quickly plateaus and remains invariant to the time window used to calculate *σ*. At higher temperatures, *σ* is dependent on the time window over which interaction energies are collected. To determine the appropriate time window in this case, we checked the movement of adjacent residues in 1 nm radius. If at least one residue crossed over the border, this would correspond to switching to a different lattice configuration and the event was noted. The frequency distribution of these times was used to obtain the time window. If the median time was less than 20 ps, the time window was an averaged value of all events that happen within the first 20 ps. As seen in Fig. S1, for polyalanine at 300 K, *σ* reaches a plateau, while at 600 K the time window is equal to about 5 ps. Fig. S2 shows *σ* values obtained by the method described above at different temperatures for C60.

We calculated *σ* in this manner for a range of temperatures, and interpolated the temperature at which *σ* crosses the theory transition line to predict the crystallization/melting temperature. For C60, the melting temperature are 405.12, 367.10, or 342.81 K depending on which one of the three all-atom force fields are used to calculate *σ*. The prediction using the united atom force field is 157.09 K. For all force fields, a single exponential interpolation function yielded a good fit over all points, although the results are insensitive to the choice of interpolation function because of close sampling near the crossing point. Because C60 begins to order within the simulation time window if the temperature is slightly below the ordering temperature, the only temperature below the ordering temperature for which we could calculate *σ* was at 100 K, for which the aggregates were trapped in the (supercooled) disordered state over the simulation window. The same procedure was repeated for PA6N9, as seen in Fig. 2b, while polypeptide calculations were all done at 300 K.

For C60, the sequence homogeneity and lack of strong interactions made it possible for us to observe spontaneous ordering of the aggregates during the course of all-atomic simulations. Melting temperatures were obtained by catching the events of crystallization during the first 25 ns of MD simulations, as we were cooling down the system, which resulted in error bars as seen in Fig. 3. In Fig. S3 we present a snapshots of AMBER99SB simulation at 400 K, in which initially disordered aggregate suddenly orders at different parts of the box. Then, the aggregate with an ordered domain was heated to 440 K. Melting was observed for all force fields except the coarse grained TraPPE-UA. To overcome the lack of spontaneous ordering for TraPPE-UA, we prepared a bipartite system consisting of 2,376 C60 polymers, with half of the chains initially ordered and the other half disordered, with the two halves separated by a domain boundary. We monitored the change in intensive and extensive parameters over time and noted the temperature at which these changes flipped sign, as seen in Fig. S4. In case of mean end-to-end distances, as well as density and volume of the systems at different temperatures, the slope changes sign between 230 K and 235 K. The observed melting temperature is therefore taken to be 230 K, which we then compared with the value of 157 K obtained by our theory.

#### Estimating *σ* for hetero-polypeptides using the additive model

After obtaining 100 ns production level trajectories from the 100 proteins listed in Table S1, we calculated the interaction energy of each type of residue with its environment. In order to get energetics that is most relevant to the bulk medium, we have excluded solvent exposed mobile residues and concentrated only in the inner cores of each protein structure. To do this, average solvent exposure and RMSD from original position are calculated for every residue in a protein, and cut off values of 0.5 nm^2^ (range 0-2.5) and 0.5 nm (range 0-1.5 nm) were used respectively for solvent exposure and RMSD in this exclusion process. Then, interaction energy (the sum of Van der Waals and Coulomb interactions) of such a buried less mobile residue and all atoms within its 1 nm proximity was calculated and averaged over the trajectory. Here, only the last 50 ns of the total 100 ns simulations was used in order to avoid any contamination from any possible initial unwanted fluctuations. Finally, distributions for all 20 amino acids were obtained as each value in these distributions corresponds to an average value over a single MD trajectory, as just outlined.

To generate *σ* values according to the additive model for random heteropolypeptides, 1000 sequences were randomly generated for a given polypeptide length, totaling 100,000 random sequences over all lengths between 2 and 101. To generate random sequences, the probability of choosing an amino acid type in a sequence was proportional to the occurrence of that amino acid within all 100 protein structures. This ensured that the generated polypeptides reflected amino acid compositions of known protein. We constructed overall polypeptide energy distribution as the superposition of the interaction energy distributions of the constituent amino acids calculated as describe above. We have made 2 important adjustment here; (i) a weighting factor is used to account for the similarity between the randomly generated sequence and each sequence of the 100 different proteins, and (ii) the means of all residue distributions were aligned prior to summation. Alignment of the mean values are needed to ensure the fixed proportion of a residue type in a given sequence. The similarity weighting factor is given by

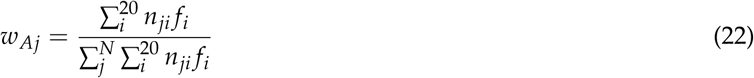

in which *w*_*Aj*_ is the weighting factor for amino acid *A* for its *j*^*th*^ occurrence in a total of 100 protein simulations. Total number of *A* in the random sequence is given by *f*_*i*_ and *n*_*ji*_ stands for total number of a given amino acid *i* around *A* in *A*’s *j*^*th*^ occurrence.

The mean energy of the overall poylpeptide is calculated by

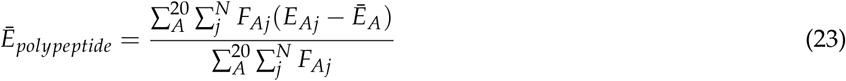

where *F*_*Aj*_ represents the weighted frequency of amino acids *A* for its *j*^*th*^ occurrence (*F*_*Aj*_ = *w*_*Aj*_*F*_*A*_), where the calculated MD interaction energy is *E*_*Aj*_. As mentioned above, each of these interaction energies are shifted by the corresponding mean value of Ē*A* to enforce the fixed amino acid composition constraint (as seen in Eq. 22).

The final standard deviation of the overall polypeptide is calculated according to:

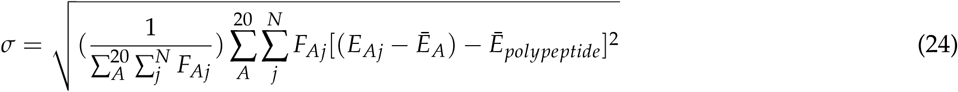

which is referred to as the *additive* model in the main text. All energy terms, as well as the final *σ*, are calculated in units of *kT*.

To estimate the error of the additive model in calculating *σ* for random hetero-polypeptide aggregates, we performed explicit atomistic calculations of fourteen different heterogeneous polypeptide sequences and calculated *σ* using the procedure described above. We then compared the results to *σ* calculated from the additive model using the amino acid interaction distributions calculated from 100 protein cores (Fig. S5). For both models, *σ* varied between 10 and 25 *kT* over a wide range of sequence complexity and composition, including the presence of charged residues. For all cases, the discrepancy was less than 5*kT*. Moreover, because the additive model underestimates *σ* except in the case of low complexity sequences, the distribution obtained by the additive model in Fig. 4b should broaden toward higher values of *σ*.

## V. Supplementary data

**Table S1:** Protein Data Bank (pdb) id numbers of the structures used to calculate amino acid interaction energy distributions.

|  |  |  |  |  |  |  |  |  |  |
| --- | --- | --- | --- | --- | --- | --- | --- | --- | --- |
| 1bw6 | 1bwi | 1ce3 | 1chv | 1gyz | 1irz | 1pc0 | 1ugl | 1vvc | 1wij |
| 1x5e | 1x9b | 1xn5 | 2a05 | 2afp | 2b3c | 2bzt | 2dad | 2dat | 2ddi |
| 2dk6 | 2dlz | 2dnu | 2e29 | 2eko | 2elk | 2gdw | 2glw | 2hgl | 2hgn |
| 2hwt | 2j8a | 2j9w | 2jzv | 2jzy | 2k22 | 2keo | 2kgf | 2kgr | 2kkg |
| 2kmc | 2kts | 2kv3 | 2l1n | 2l4m | 2l6q | 2l92 | 2l93 | 2lea | 2lgh |
| 2lgx | 2lnm | 2lu2 | 2lvf | 2m3a | 2m4v | 2m8r | 2m94 | 2mah | 2mmu |
| 2mry | 2n2q | 2n2r | 2n2z | 2n3l | 2n4o | 2n69 | 2n81 | 2n98 | 2na2 |
| 2nar | 2o2w | 2rq1 | 2ru1 | 2ruh | 2uz5 | 2v1n | 2vc8 | 2x43 | 2ydl |
| 2yqd | 2ys9 | 3c4s | 3jte | 3zg4 | 4bwh | 4d4w | 4fpr | 4hqa | 4htw |
| 5f6a | 5gvq | 5iaz | 5jfb | 5k57 | 5k6d | 5kpe | 5lxl | 5mfy | 5ttt |

**Figure S1:**
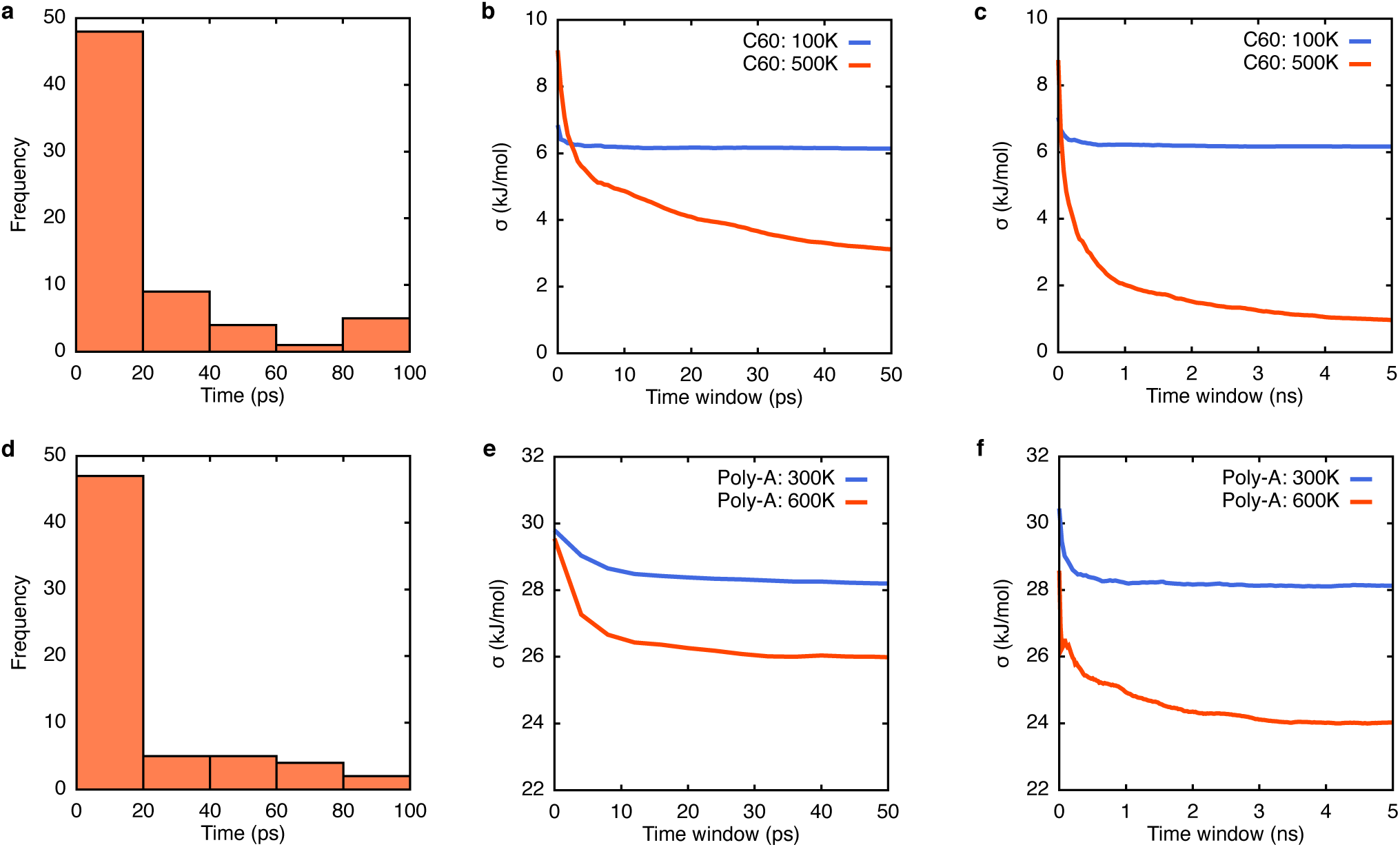
Protocol for calculating standard deviation of interaction energy for bulk media. Frequencies of residues leaving the 1 ns radius around a residue of interest at particular times for C60 (a) and polyalanine (d). Changes in σ averaged over time windows from 0 to 100 ps for C60 (b) and polyalanine (e), as well as from 0 to 5 ns for C60 (c) and polyalanine (f).

**Figure S2:**
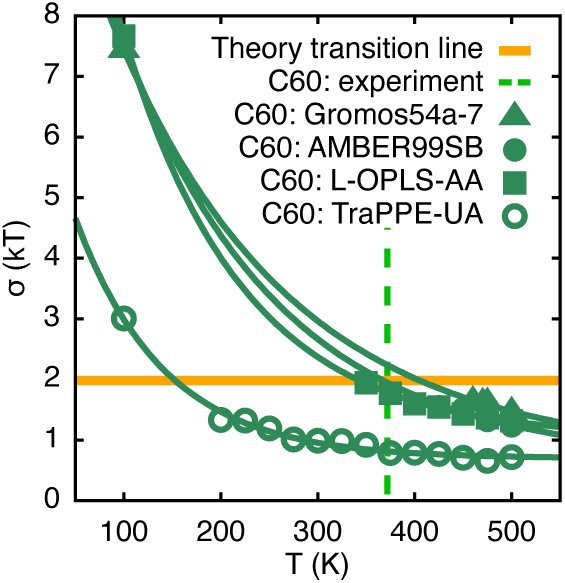
Theoretical transition points prediction for C60 using different force fields with experimental transition temperature as a dashed line.

**Figure S3:**
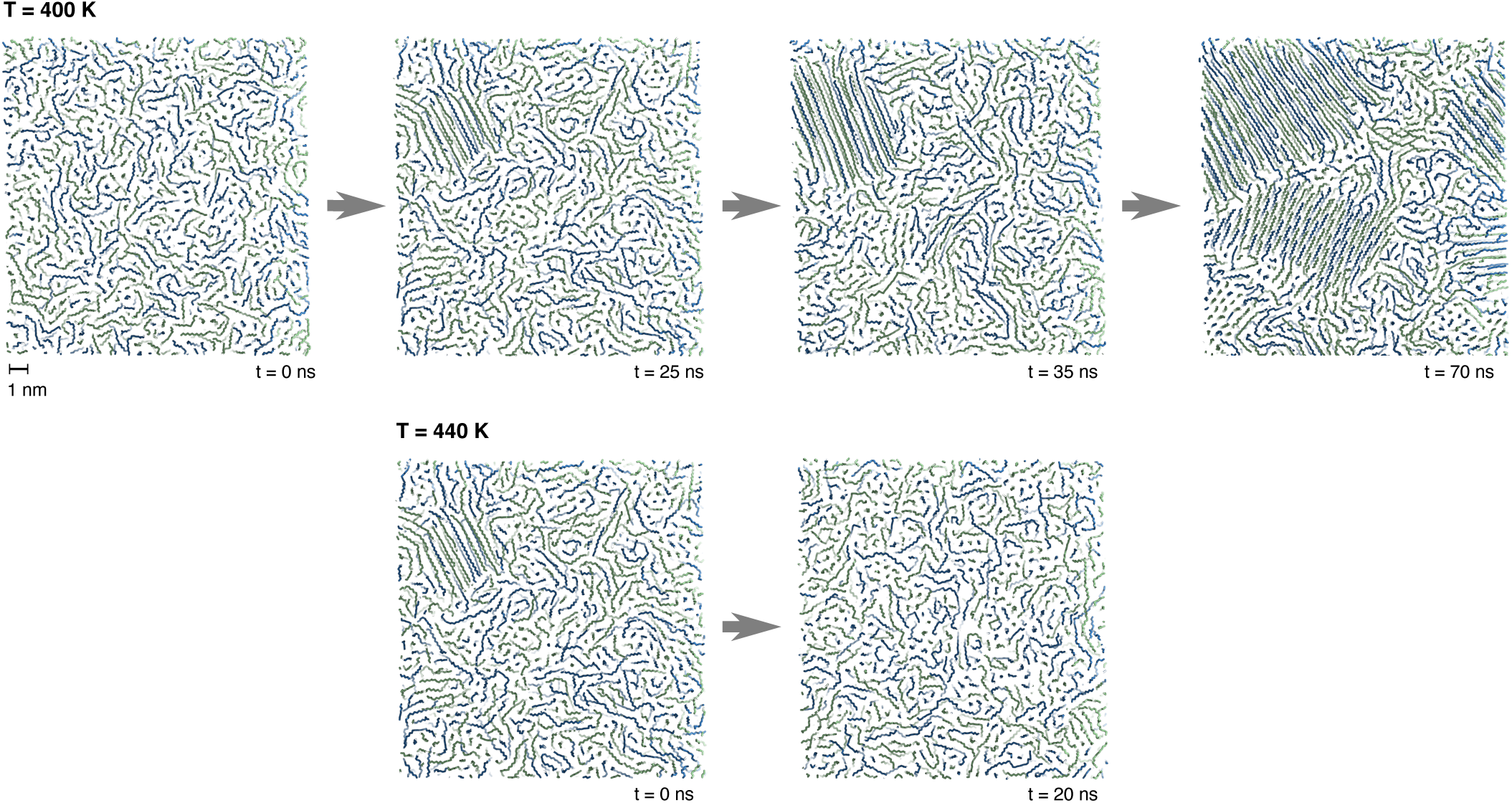
Observed spontaneous ordering of C60 using AMBER99SB force field at 400 K (0, 25, 35, 70 ns), as well as melting of the ordered domain at 440 K after 20 ns.

**Figure S4:**
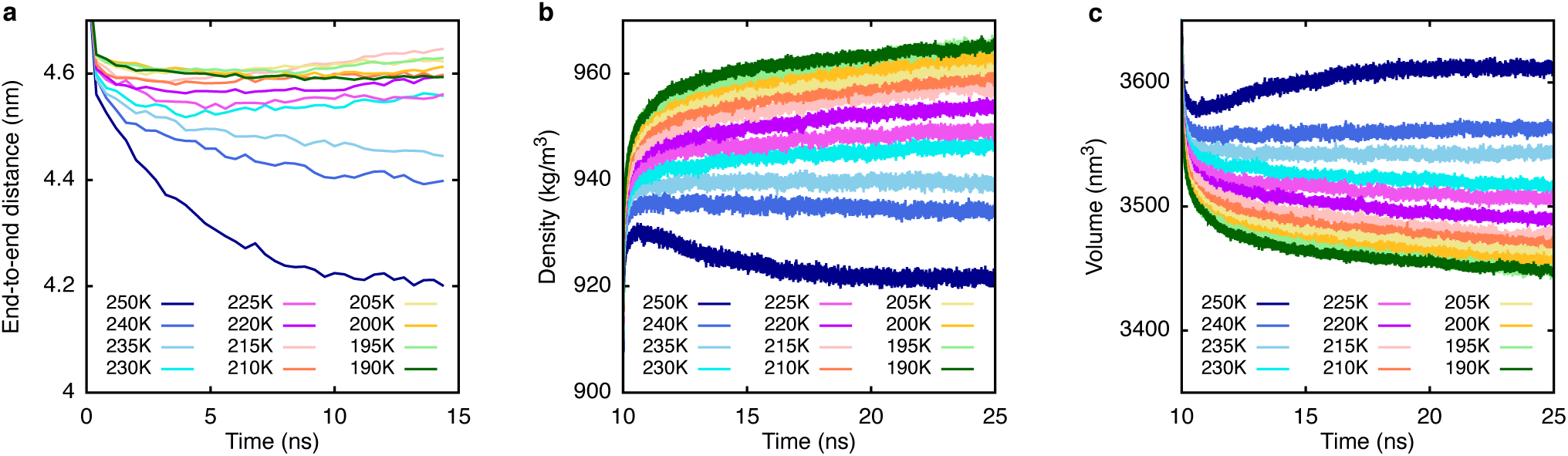
Change in C60 intensive and extensive variables using TraPPE-UA (united atom) force field at different temperatures. a, Changes in mean end-to-end distances of all polymers with time. b, Changes in density of the systems beginning at 10 ns of simulations. c, Changes in volume of the systems beginning at 10 ns of simulations.

**Table S2:**
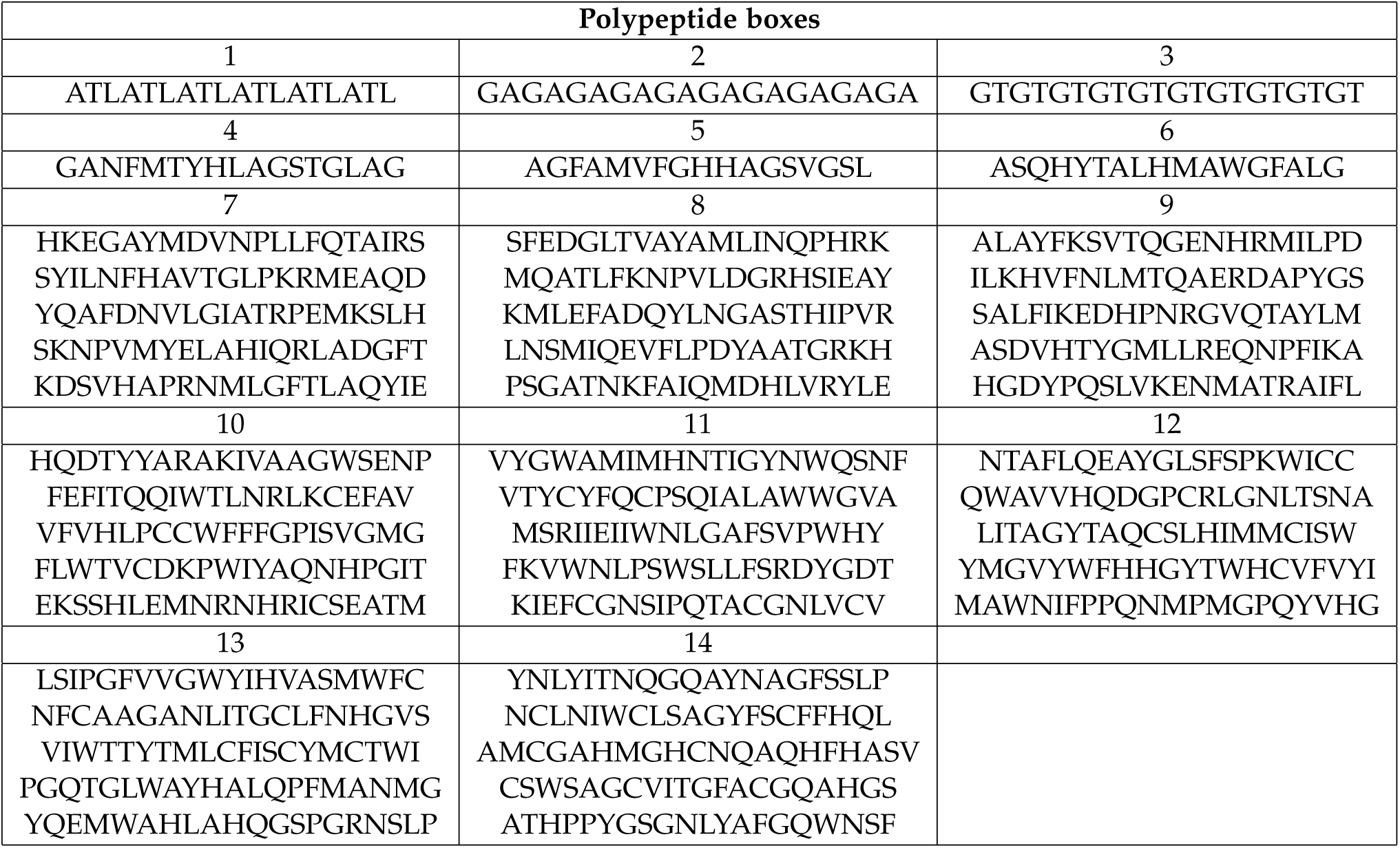
Composition of boxes containing heterogenous polypeptide chains used for explicit bulk media MD simulations.

**Figure S5:**
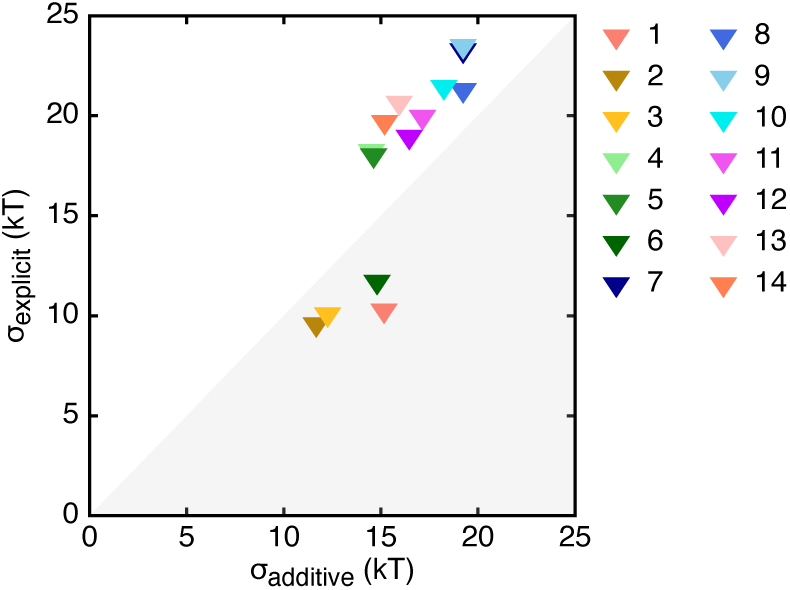
Comparison of standard deviations of interaction energies of polypeptides calculated using the quick additive method (x axis) and explicit (bulk media) molecular dynamic simulations (y axis). Boxes’ composition can be seen in Table S2.

